# Identification of novel differentially methylated sites with potential as clinical predictors of impaired respiratory function and COPD

**DOI:** 10.1101/473629

**Authors:** Mairead L Bermingham, Rosie M Walker, Riccardo E. Marioni, Stewart M Morris, Konrad Rawlik, Yanni Zeng, Archie Campbell, Paul Redmond, Heather C Whalley, Mark J Adams, Prof. Caroline Hayward, Prof. Ian J Deary, Prof. David J Porteous, Prof. Andrew M McIntosh, Kathryn L Evans

**Affiliations:** Centre for Genomic and Experimental Medicine, Institute of Genetics and Molecular Medicine, University of Edinburgh, Edinburgh, UK; Centre for Cognitive Ageing and Cognitive Epidemiology, University of Edinburgh, Edinburgh, UK; Division of Genetics and Genomics, The Roslin Institute and Royal (Dick) School of Veterinary Studies, University of Edinburgh, Easter Bush, Roslin, UK; Medical Research Council Human Genetics Unit, Institute of Genetics and Molecular Medicine, University of Edinburgh, Edinburgh, UK; Usher Institute of Population Health Sciences and Informatics, University of Edinburgh, Edinburgh UK; Division of Psychiatry, University of Edinburgh, Royal Edinburgh Hospital, Edinburgh, UK

## Abstract

**Background:** The causes of poor respiratory function and COPD are incompletely understood, but it is clear that genes and the environment play a role. As DNA methylation is under both genetic and environmental control, we hypothesised that investigation of differential methylation associated with these phenotypes would permit mechanistic insights, and improve prediction of COPD. We investigated genome-wide differential DNA methylation patterns using the recently released 850K Illumina EPIC array in the largest single population sample to date.

**Methods:** Epigenome-wide association studies (EWASs) of respiratory function and COPD were performed in peripheral blood samples from the Generation Scotland: Scottish Family Health Study (GS:SFHS) cohort (N=3,791; 274 COPD cases and 2,928 controls). In independent COPD incidence data (N=150), significantly differentially methylated sites (DMSs; p<3.6×10^−8^) were evaluated for their added predictive power when added to a model including clinical variables, age, sex, height and smoking history using receiver operating characteristic analysis. The Lothian Birth Cohort 1936 (LBC1936) was used to replicate association (N=895) and prediction (N=178) results.

**Findings:** We identified 29 respiratory function and/or COPD associated DMSs, which mapped to genes involved in alternative splicing, JAK-STAT signalling, and axon guidance. In prediction analyses, we observed significant improvement in discrimination between COPD cases and controls (p<0.05) in independent GS:SFHS (p=0.014) and LBC1936 (p=0.018) datasets by adding DMSs to a clinical model.

**Interpretation:** Identification of novel DMSs has provided insight into the molecular mechanisms regulating respiratory function and aided prediction of COPD risk.

**Funding:** Wellcome Trust Strategic Award 10436/Z/14/Z.

**Research in context:** *Evidence before this study:* We searched for articles in PubMed published in English up to July 25, 2018, with the search terms “DNA methylation” and “respiratory function”, or “COPD”. We found some evidence for association between differential DNA methylation and both respiratory function and COPD. Of the twelve previous studies identified, eight used peripheral blood samples (sample size [N] range = 100-1,085) and four used lung tissue samples (N range = 24-160). The number of CpG loci analysed range from 27,578 to 485,512. These studies have not identified consistent changes in methylation, most likely due to a combination of factors including small sample sizes, technical issues, phenotypic definitions, and study design. In addition, no previous study has: analysed a sample from a large single cohort; used the recently released Illumina EPIC array (which assesses ~850,000 CpG loci); adjusted methylation data and phenotype for smoking history, or used both prevalent and incident COPD electronic health record data.

*Added value of this study:* To our knowledge, this is the largest single cohort epigenome-wide association study (EWAS) of respiratory function and COPD to date (N=3,791). After applying stringent genome-wide significance criteria (P <3.6×10^−8^), we found that DNA methylation levels at 29 CpG sites in peripheral blood were associated with respiratory function or COPD. Of these 29, seven were testable in an independent population sample: all seven showed consistent direction of effect between the two samples and three showed replication (p<0.007 [0.05/7 CpG sites tested]). Our results suggest that adjustment of both the phenotypic and the DNA methylation probe data for smoking history, which has not been carried out in previous studies, reduces the confounding effects of smoking, identifies larger numbers of associations, and reduces the heterogeneity of effects across smoking strata. We used gene set enrichment and pathway analyses, together with an approach that combines DNA methylation results with gene expression data to provide evidence for enrichment of differentially methylated sites in genes linked to alternative splicing, and JAK-STAT signalling and axon guidance. Finally, we demonstrated that the inclusion of DNA methylation data improves COPD risk prediction over established clinical variables alone in two independent datasets.

*Implications of all the available evidence:* There is now accumulating evidence that DNA methylation in peripheral blood is associated with respiratory function and COPD. Our study has shown that DNA methylation levels at 29 CpG sites are robustly associated with respiratory function and COPD, provide mechanistic insights, and can improve prediction of COPD risk. Further studies are warranted to improve understanding of the aetiology of COPD and to assess the utility of DNA methylation profiling in the clinical management of this condition.

## Introduction

Respiratory function is influenced by both environmental factors and genetic factors, with heritability estimates ranging from 39 to 66%.^1,2^ Epigenetic modifications are at the interface of genetics and the environment. DNA methylation, the covalent binding of a methyl group to the 5′ carbon of cytosine-phosphate-guanine (CpG) dinucleotide sequences in the genome, is an epigenetic modification of DNA that is associated with gene expression. The results of previous epigenome-wide association studies (EWASs) of spirometric measures of respiratory function and respiratory disease have produced inconsistent results, with some identifying significant associations,^3–6^ and others not.^7–9^ Moreover, there has been little consistency between the positive findings reported.^9,10^

Studies of lung tissue^5,8^ have been constrained by sample availability, with the largest study to date comprising 160 subjects.^5^ Inconsistency amongst the results of the peripheral blood-based studies^3,6,7,9,11^ is likely to be due to a number of factors, including small sample size (e.g., two studies had less than 200 samples)^6,11^ and/or investigation of a relatively small number (~27,000) of CpG loci.^3,11^ The study with the largest number of samples (n=1,085) analysed only 27,000 CpG loci, while the largest study using the 450K array (the predecessor to the array used here) analysed 920 samples.^12^ Differences in spirometric measures, definitions of COPD, study population characteristics and study design, in particular in the method used to adjust for smoking history, are also likely to be important sources of variation.^9,10^

Here we sought to identify robust associations by assessing methylation in a large single cohort sample, applying a more rigorous correction for smoking history and by performing sensitivity analyses. In contrast to prior studies, we used the recently released Illumina EPIC array, which interrogates over 850,000 methylation sites. All 3,791 individuals in our sample were from a single cohort with extensive and consistent phenotyping comprising clinical investigation, questionnaire, and linkage to routine medical health records. The cross-sectional design of prior studies has limited their capacity to distinguish cause and effect.^**10**^ We mitigated this issue by testing our findings for their predictive power in an independent subpopulation of 150 participants with incident COPD who were free of disease at the time of blood sampling. Finally, where data were available, we attempted to replicate our EWAS and prediction findings in an independent cohort, LBC1936, drawn from the same population.

## Material and methods

A flow chart showing the overall study design is outlined in figure 1, and full description of the methods is provided in the appendix.

**Figure 1.**
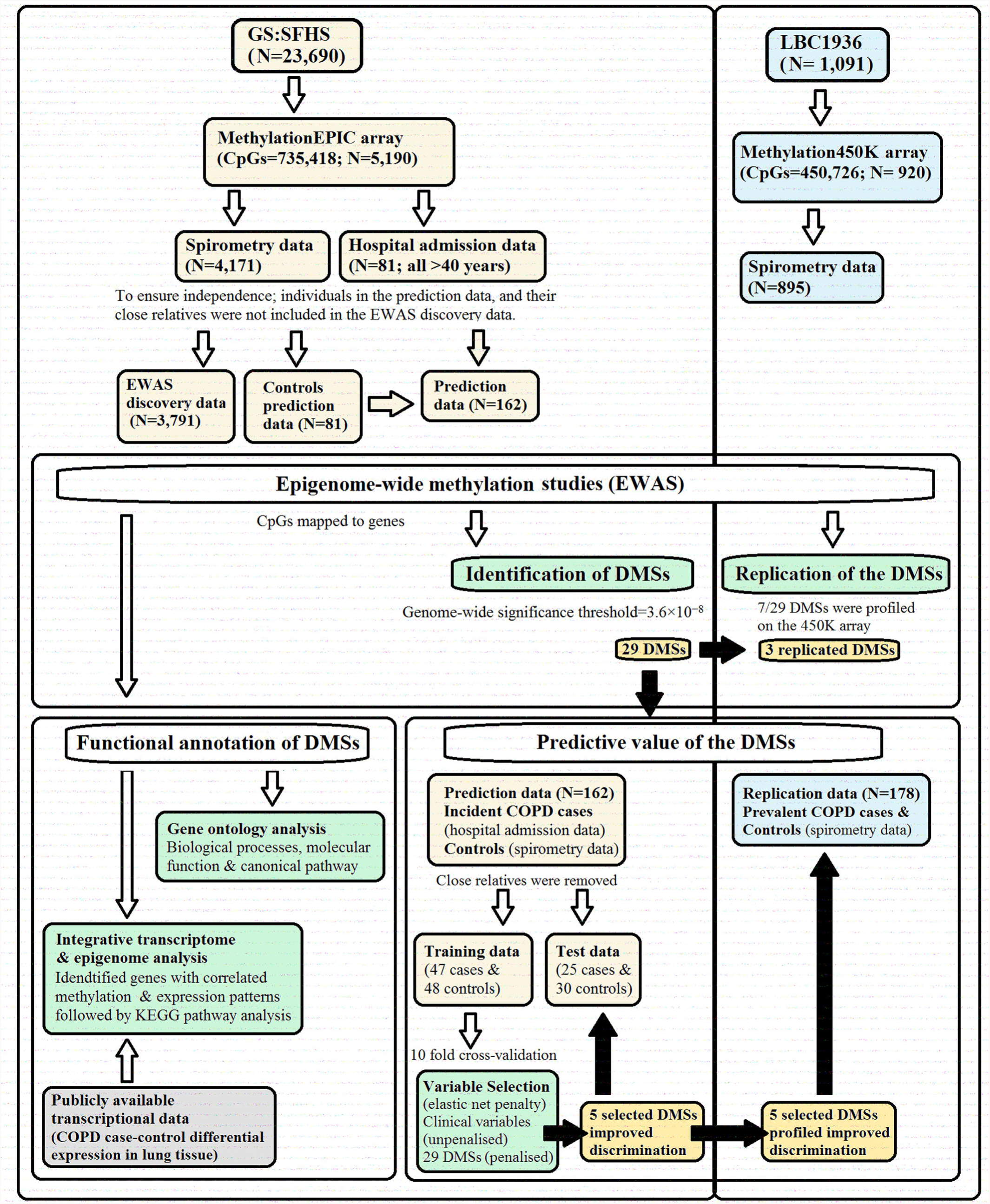
Flow-chart showing the analysis pipeline. Direction of the arrows represents the workflow of the study design with performed analysis indicated. Lemon and blue boxes represent the in the discovery Generation Scotland: Scottish Family health study cohort and the replication Lothian Birth Cohort of 1936 (LBC1936) data sets respectively. The grey box indicates input data of COPD case-control differential expression in lung tissue. The green boxes indicate the analyses undertaken. The black arrows and gold boxes indicate output of significant results.

### Epigenome-wide association study

#### Cohort information

The Generation Scotland Scottish Family Health Study (GS:SFHS) and Lothian Birth Cohort of 1936 (LBC1936) comprise 23,690 and 1,091 participants, respectively. Medical Research Ethics was obtained for all components of GS:SFHS and LBC1936. Written informed consent was obtained from all participants.

#### Genome‐wide methylation profiling

In the GS:SFHS cohort, DNA methylation data was obtained from 5,190 participants using peripheral blood collected at baseline.^13^ DNA methylation was assessed using the Infinium MethylationEPIC BeadChip. Quality control procedures were implemented to identify and remove unreliable probes and samples, and probes on the X and Y chromosomes were excluded leaving data for 735,418 methylation loci in 5,190 individuals. In the LBC1936, DNA methylation was assessed in whole blood samples from 1,004 participants using the Illumina HumanMethylation450 BeadChip. Low-quality probes and samples, and probes on the X and Y chromosomes were removed, leaving 450,726 probes and 920 samples for inclusion in the analysis. M-values were calculated for both datasets. The M-values were then pre-corrected for relatedness (in GS:SFHS), array processing batch and estimated cell counts.

#### Trait data

Respiratory function was assessed at the time of blood sampling in 4,193 GS:SFHS and 895 LBC1936 participants with methylation data. Forced expiratory volume in one second (FEV_1_) and forced vital capacity (FVC) were measured in litres, using spirometry. Quality control of the phenotype data was undertaken to exclude participants with inaccurate spirometry or covariate data; 4,171 and 895 individuals were retained in the GS:SFHS and LBC1936 samples respectively.

Following the Global Initiative for Obstructive Lung Disease (GOLD) criteria, participants with predicted FEV_1_ (from NHANES III equations) <80% and FEV_1_/FVC <0.7 were classified as COPD case subjects.^14^ Individuals with FEV_1_ >80% predicted and FEV_1_/FVC>0.7 were classified as control subjects.

The GS:SFHS dataset (N=4,171) was then divided into an incident COPD dataset (i.e., where COPD developed after recruitment to the cohort) for prediction analysis and a discovery EWAS dataset. The prediction dataset (described below) comprised of incident COPD cases and matched controls. To ensure independence, individuals in the prediction sample and their close relatives (identity by state [IBS]>0.05) were not included in the discovery dataset; leaving 3,791 for inclusion in the EWAS analysis.

#### Identification of differentially methylated sites (DMSs)

FEV1, FVC, FEV1/FVC, and COPD phenotypic data were pre-corrected for age, age^2^, sex, height, height^2^, smoking status (current smoker, former smoker [< 12 months], former smoker [≥ 12 months], and never smoked), and pack-years. FVC data was additionally pre-corrected for weight. Linear regression models were run in the *limma* package in R, fitting each CpG site (corrected M-values) as the dependent variable, and respiratory function traits or COPD, age, sex, smoking status, pack-years and the first 20 principal components from the corrected M-values. The genome-wide significance threshold was set at 3.6×10^−8^.^15^

#### Sensitivity analyses

To assess the impact of pre-correction of the traits for smoking history, we undertook sensitivity analyses, in which FEV1, FVC, FEV1/FVC, and COPD were not pre-corrected for smoking status and pack-years. To assess the generalisability of the estimated effects in older adults and according to smoking status (ever smokers and non-smokers), the data was censored by age (>40 years; an age group with greater risk of COPD^14^) and stratified by smoking status. For each set of analyses, we conducted separate random-effects meta-analyses using the *metafor* package in R for each trait-specific genome-wide significant DMS.

### Probe annotation and epigenetic regulation of gene expression

DNA methylation probes were mapped to genes based on the IlluminaHumanMethylationEPICanno.ilm10b2.hg19 library. For each trait, methylation probes were filtered at p<0.001, and probes that mapped to genes extracted. For biological processes and molecular function, and canonical pathway enrichment analyses, DMSs were analyzed in the Database for Annotation, Visualization and Integrated Discovery (DAVID) database and Ingenuity Pathway Analysis (IPA) software respectively.

The Benjamini Hochberg, False Discovery Rate method, was used to correct for multiple-testing with *p*<0.05 being considered significant. We used the *Significance-based Modules Integrating the Transcriptome and Epigenome* (SMITE) package in R to combine summary statistics from publicly available COPD lung gene expression data^16^ with methylation results from this study. Trait-specific gene modules (set of genes with shared regulation; p<0.05 and 10–500 genes) were then identified and subjected to KEGG pathway enrichment analysis and terms with a p<0.05 were held as significant.

### Prediction

#### Case-control data

Incident COPD cases in GS:SFHS were defined as any hospital admission where the primary diagnosis was assigned an ICD-10 J40 to J44 code. During follow-up, 81 GS:SFHS participants (all 40 years or older) with DNA methylation data developed COPD. To obtain a balanced dataset for model training an equal number of controls ≥40 years of age were selected at random from those with no-self report, spirometry-defined, or ICD-10 diagnosis of COPD. Participants with missing records and closely related individuals (IBS>0.05) were excluded; leaving 72 COPD cases and 78 controls. The data was then separated into a training set of 47 COPD cases and 48 controls, and a test set of 25 COPD cases and 30 controls.

As hospital admission data were not available for the LBC1936 cohort, spirometry data were used to define case-control status. In total, 89 participants with DNA methylation data had prevalent COPD [GOLD stage ≥2 cases]. Controls were selected at random from the participants with DNA methylation data in a 1:1 ratio to case participants. This dataset was used to replicate the prediction findings from GS:SFHS.

#### Model selection

For the training data, the reduced model, including clinical risk factors, age, age^2^, sex, height, height^2^, smoking status (current, former and never), and pack-years of smoking^6^ was constructed using unpenalized logistic regression. The full model, including DMSs and clinical risk factors, was constructed using penalized logistic regression with an elastic net penalty. Selection of the full model was conducted based on 10-fold cross-validation (appendix p28) using the R package *caret*. The optimal model was selected based on the maximum mean area under the curve (AUC). Final models were constructed using the complete training set and evaluated on the independent test and replication datasets.

#### Model evaluation

Comparison of the predictive performance of the models was carried out using the AUC in the *pROC* R package. The incremental value of the DMS to predict COPD risk, when added to the model with established clinical predictors was assessed using the integrated discrimination improvement (IDI), and binary net reclassification improvement (NRI) measure. Finally, we performed decision curve analysis to estimate the potential clinical usefulness of the models in the ‘rmda’ R package.

## Results

### EWAS sample characteristics

The discovery sample for respiratory function traits comprised 3,791 individuals from GS:SFHS. For the COPD analysis, there were 274 cases and 2,928 controls (table 1; figure 1).

**Table 1.** Characteristics of Generation Scotland: Scottish Family Health Survey (GS:SFHS) participants (n= 3,791) in the epigenome-wide association study discovery population.

| Characteristics | COPD cases<br>(n = 274) | Controls<br>(n = 2,928) | GOLD stage 1<br>(n=589) |
| --- | --- | --- | --- |
| **Age, years** | 53.96 ± 13.59 | 46.97 ± 13.37 | 52.16 ± 12.74 |
| **Sex:** |  |  |  |
| – Male | 93 (33.9) | 1,167 (39.9) | 222 (37.7) |
| – Female | 181 (66.1) | 1,761 (60.1) | 367 (62.3) |
| **Height, cm** | 166.39 ± 8.76 | 167.85 ± 9.08 | 166.96 ± 9.48 |
| **Weight, kg** | 73.49 ± 15.63 | 75.82 ± 16.32 | 76.57 ± 16.87 |
| **Smoking status:** |  |  |  |
| – Never | 96 (35.0) | 1,609 (55.0) | 250 (42.5) |
| – Former (> 12 months) | 74 (27.0) | 716 (24.4) | 164 (27.8) |
| – Former (< 12 months) | 3 (1.1) | 88 (3.0) | 20 (3.4) |
| – Current | 89 (32.5) | 445 (15.2) | 133 (22.6) |
| – Missing records | 12 (4.4) | 70 (2.4) | 22 (3.7) |
| **Pack-year:** |  |  |  |
| – Former smokers (> 12 months) | 25.47 ± 31.84 | 16.67 ± 20.30 | 22.83 ± 22.87 |
| – Former smokers (< 12 months) | 30.00 ± 31.05 | 15.16 ± 15.75 | 18.35 ± 20.20 |
| – Current smokers | 22.97 ± 19.15 | 15.65 ± 15.94 | 23.59 ± 16.85 |
| **Lung function:** |  |  |  |
| – FEV<sub>1</sub>, litres/second | 2.01 ± 0.60 | 3.24 ± 0.76 | 2.58 ± 0.66 |
| – FVC, litres/second | 3.45 ± 0.93 | 4.07 ± 0.94 | 3.72 ± 1.08 |
| – FEV<sub>1</sub>/FVC | 0.59 ± 0.10 | 0.80 ± 0.06 | 0.71 ± 0.08 |
| – FEV<sub>1</sub> percent predicted | 66.96 ± 11.44 | 99.56 ± 11.16 | 83.97 ± 12.98 |
| – FVC percent predicted | 98.90 ± 13.09 | 99.49 ± 11.90 | 94.92 ± 20.80 |

**Abbreviations:** COPD, Chronic obstructive pulmonary disease; FEV<sub>1</sub>, Forced expiratory volume in one second; FVC, Forced vital capacity. Figures shown are the mean ± standard deviation or n (%).

### Differentially methylated sites

EWASs for the three respiratory function traits (FEV1, FVC and FEV1/FVC) and COPD on the discovery data identified 30 genome-wide significant associations (p<3.6×10^−8^; figure 2; table 2; appendix p6-7), representing 29 DMSs from 26 annotated genes. We found only marginal evidence of genomic inflation (max=1.13) across traits (appendix p6 & 29). Eleven of the 26 genes that contain DMSs have previously been implicated by genetic, EWAS and functional analysis in respiratory function or disease (excluding cancer; appendix p8-9). Three FEV_1_ related-DMSs mapped to the SOCS3 gene (table 2); DNA methylation levels at these sites are highly correlated (appendix p30).

**Table 2.**
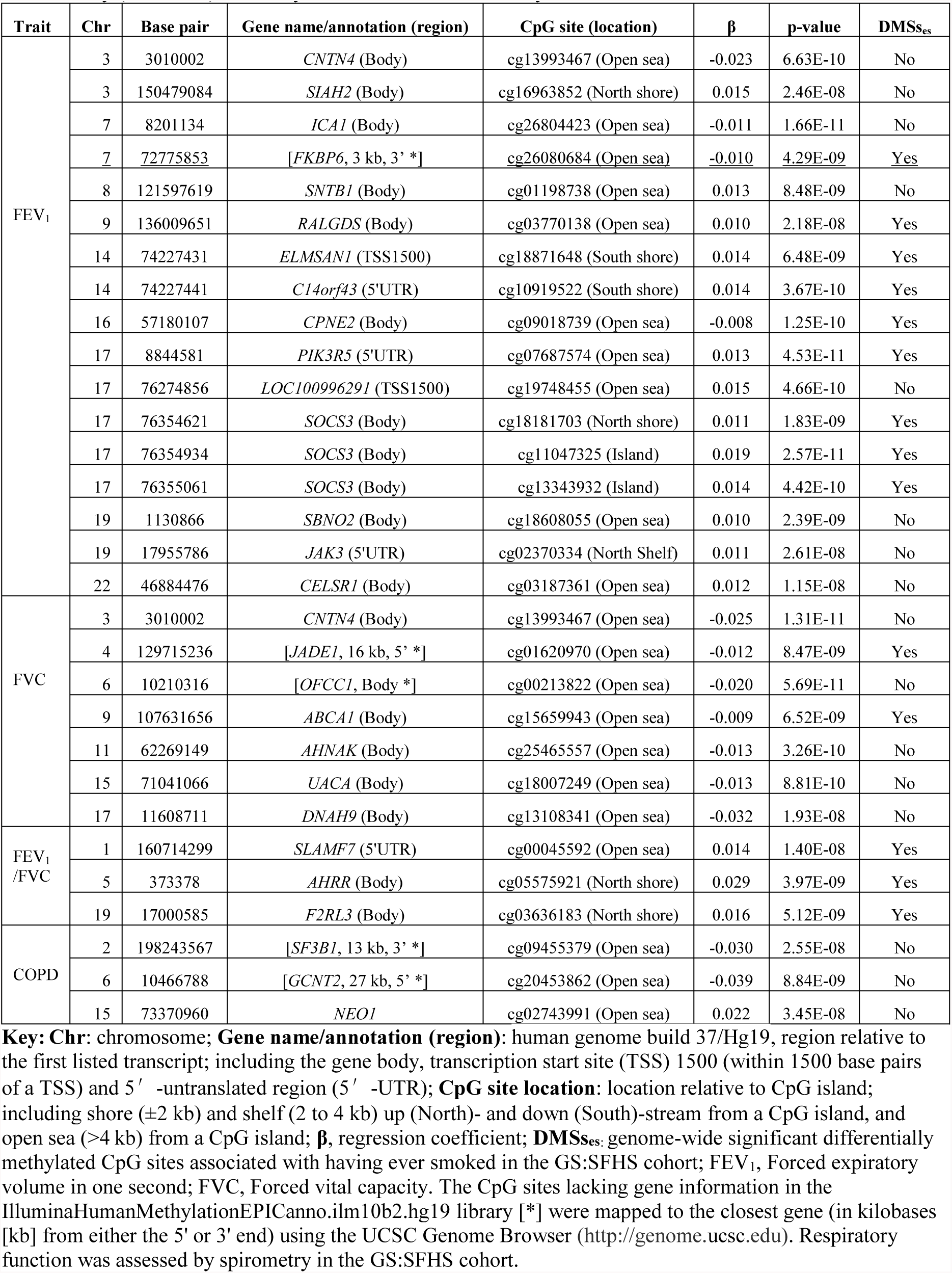
Genome-wide significant differentially methylated sites (DMSs) associated with the respiratory function traits or chronic obstructive pulmonary disease (COPD) in the Generation Scotland Scottish Family Health Study (GS:SFHS) discovery data. Results are ordered by chromosomal location.

**Figure 2.**
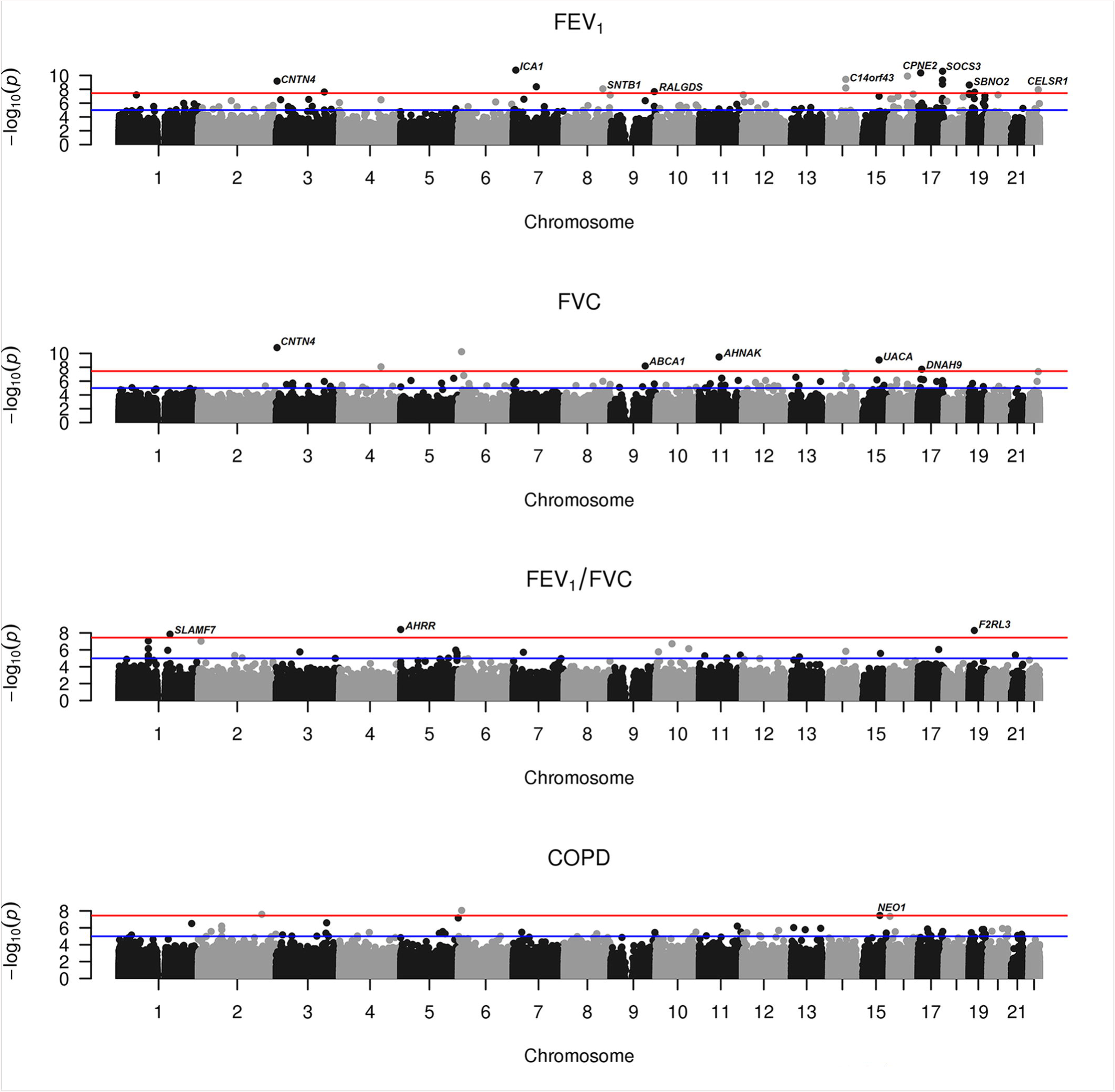
Manhattan plots of epigenome-wide association results for FEV_1_ (forced expired volume in 1 s), FVC (forced vital capacity; bottom), FEV_1_/FVC and COPD (chronic obstructive pulmonary disease) from the discovery Generation Scotland: Scottish Family health study cohort data. The red line correspond to the genome-wide (*P* = 3.6 × 10^−8^) and suggestive (*P* = 1.0 × 10^−5^) significance level. Labels are for the nearest gene to genome-wide significant CpG sites.

We next attempted to replicate these findings in 895 individuals from the LBC1936 (appendix p10). No other Illumina EPIC dataset was available, but seven of the 29 DMSs identified in the discovery dataset had been profiled using the HumanMethylation450 BeadChip, which had been applied to the LBC1936. Of these seven, two FEV_1_-associated DMSs (cg18181703 in *SOCS3* and cg18608055 in *SBNO2*) and one FEV1/FVC ratio-associated DMS (cg03636183 in *F2RL3*) replicated in LBC1936 (Bonferroni-corrected p≤0.00714). For all seven probes, however, the direction of the effects were the same in the two datasets (appendix p11).

### Sensitivity analyses

As age and smoking affect both DNA methylation and lung function,^17,18^ we undertook sensitivity analyses for each significant DMS for each trait. The associations between each significant DMS and its associated phenotype were assessed in older adults (>40) and smoking strata (ever smokers and non-smokers). Meta-analysis was used to compare regression coefficients from the discovery dataset and the sensitivity analyses. All but three of the associations identified in the discovery dataset were robust to differences in age and smoking status (p<0.05/30 DMSs; appendix p12-13). The association between FVC and cg00213822 in *OFCC1* was primarily driven by non-smokers, whereas, the associations with FEV1/FVC and the established smoking-associated DMSs, cg05575921, in *AHHR*, and cg03636183, in *F2RL3*, were primarily driven by smokers (appendix p31).

We carried out sensitivity analyses in which the trait data were not pre-corrected for smoking history (appendix p14-17). Three of the identified associations were affected by pre-correction of the traits (appendix p17 & 32). In addition, an FVC-related DMS was identified at cg03187361 only when the trait data was not pre-corrected for smoking history. Pre-correction for smoking history substantially reduced the heterogeneity of the effect size estimates of the association between COPD and DNA methylation at cg09455379 and cg02743991 across the age and smoking strata. This strengthened associations, which attained genome-wide significance.

### Gene ontology analysis

For each trait, to explore whether genes with DMSs share functional features, we filtered methylation probes at p<0.001 and performed biological processes (appendix p33), molecular function (appendix p34) and canonical pathway enrichment analyses (appendix p18-25). In the study of molecular functions, we found that each of the four traits were significantly enriched for genes linked to the alternative splicing and phosphoprotein categories (appendix p34), while DMSs associated with FEV_1_ and COPD showed enrichment for “cytoplasm organization and biogenesis”, and “disease mutation” respectively (appendix p34). Many of the canonical pathways identified were related to signalling, including apoptosis (appendix p18), cardiovascular signalling (appendix p20) and neurotransmission (appendix p25).

### Integrative analysis of methylation and expression

To investigate the functional relevance of the methylation changes, we integrated transcriptional (COPD case-control differential expression in lung tissue)^16^ and epigenetic (from this study) datasets to identify functional gene modules for the traits under study. This analysis identified two significant modules (p<0.05) containing 27 and 34 genes with correlated differential methylation associated with FEV_1_ and expression in COPD, respectively (appendix p26, p35-36). DMSs mapped to two genes *SIAH2* and *NEO1* in module 1 (appendix p35) and *SOCS3* in module 10 (appendix p36). Gene enrichment analysis revealed that the top pathway for module 1 was axon guidance, while the top pathways for module 10 were cytokine-cytokine receptor interaction and JAK-STAT signalling (p<0.05; appendix p26). No significant functional gene modules were identified for the other phenotypes.

### Predictive value of the DMS

To determine the predictive value of DMSs in the prognosis of COPD, we used an independent training and test set design to predict COPD risk in GS:SFHS and LBC1936. We calculated the improvement in prediction quality of a model where genome-wide significant DMSs with all traits were added to the reduced model, which included the clinical variables: age, age^2^, sex, height, height^2^, smoking status and pack-years of smoking^6^. For descriptive statistics of the prediction datasets see appendix p27. Discrimination of the full model in the GS:SFHS test data was excellent (AUC=0.923[95% CI: 0.847-0.998]; appendix p37) and calibration was good (Hosmer–Lemeshow goodness-of-fit p=0.368; appendix p38). Addition of DMSs to the reduced model led to a modest improvement in the AUC of 0.021 (95%CI:0.012-0.030; p=0.171), and resulted in a significant improvement in both discrimination (IDI: 0.065[95%CI: 0.013-0.117; p=0.014]; appendix p39) and reclassification (NRI: 0.153[0.107-0.296, p=0.035]; figure 3).

**Figure 3.**
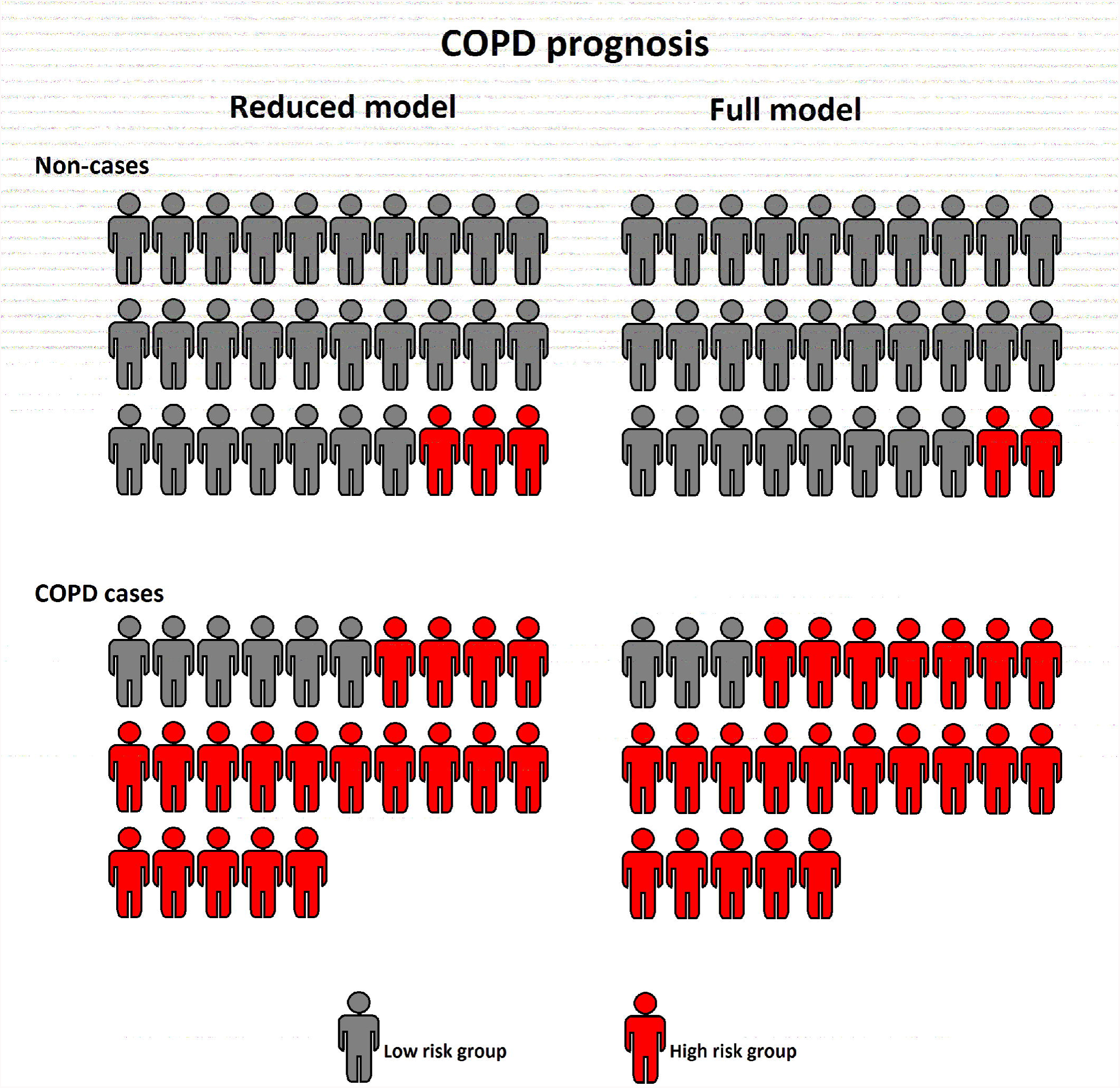
Predicted risk groups in the independent Generation Scotland: Scottish Family health study (GS:SFHS) test data (30 non-cases and 25 COPD cases) were defined as: low-risk group < 60% and higher-risk group ≥ 60%. The risk score from the reduced model built in the training data from the discovery GS:SFHS cohort contains information on sex, age, age^2^, height, height^2^, smoking status [never, former and current smoker] and pack-years of smoking. The risk score from the full model contains information from the reduced model and differentially methylated sites associated with respiratory function and prevalent chronic obstructive pulmonary disease (COPD) in the discovery GS:SFHS data. Grey non-cases and red COPD cases denote correctly classified individuals.

We examined glmnet’s variable importance measures to determine which DMSs contributed most to the increased discriminatory power. Five DMSs: cg03636183 (*F2RL3*); cg03770138 (*RALGDS*), cg18181703 (*SOCS3*), cg01620970 (*JADE1*) and cg16963852 (*SIAH2*); were retained for prediction (appendix p40).

We next assessed the full model in the LBC1936 replication sample, which comprised 89 cases and 89 controls. Due to differences in array coverage, only two of the DMSs retained in the full model built in the GS:SFHS training data could be tested in LBC1936 (cg03636183 and cg18181703). Addition of the two sites to the reduced model led to a significant improvement in the AUC of 0.030(95%CI: 0.025-0.035; p=0.029), and discriminative power (IDI: 0.039[95%CI: 0.006-0.067; p=0.018]) and total reclassification (NRI: 0.135[0.043-0.227, p=0.004])

Decision curve analysis showed that the model incorporating the DMSs had good clinical applicability and was superior to the reduced model over a wide range of threshold probabilities in the discovery and replication data (appendix p 41 & 42).

## Discussion

We performed EWASs of three respiratory function traits and COPD in DNA extracted from peripheral blood using the high-density Illumina EPIC array, in 3,471 individuals from a single cohort. These analyses identified 29 DMSs (28 novel), of which 26 are associated with respiratory function and three with COPD in the discovery GS:SFHS data. Data were available to test seven of the 29 DMSs for replication in an independent dataset; three associations replicated. Adjustment of both the phenotypic and DNA methylation data for smoking history appeared to reduce the confounding effects of smoking, identify more associations, and reduce the heterogeneity of effect estimates across smoking strata. Incorporation of a subset of the identified DMSs into a model composed of established clinical variables improved discrimination of individuals at-risk of COPD in two independent samples. Finally, functional annotation provided insights into the biology of these phenotypes.

Eleven of the 26 genes that contain DMSs have been previously linked to respiratory function or disease (appendix p8-9). In four cases, these links come from studies in lung tissue: DNA methylation changes in *ABCA1* in lung tissue has been reported to be associated with pulmonary arterial hypertension; differential expression of *ABCA1* and *DNAH9* has been reported in lung tissue of patients with COPD and primary ciliary dyskinesia respectively and pathological changes in lung tissue have been reported following knockdown and knockout of *SLAMF7*, *ABCA1*, and *SOCS3*.

Three DMSs showed replication. The first, cg18181703, is one of three FEV_1_-associated DMS in *SOCS3*, which modulates the lung inflammatory response,^19^ and JAK-STAT signal transduction.^20^ Transcriptional down-regulation of *SOCS3* has been observed in COPD patients.^21^ Differentially methylated sites in *SOCS3* within a FEV_1_-related gene module in this study were correlated with differential gene expression in lung tissue of COPD patients.^16^ A DMS in *SOCS3* was the third most predictive DMS in the prognostic model for COPD in the GS:SFHS cohort. Inclusion of this DMS in the prediction model also improved the prediction of prevalent COPD risk in the LBC1936.

The second replicated DMS, cg18608055, is a FEV_1_-associated DMS in *SBNO2*, which encodes a transcriptional co-regulator of the pro-inflammatory cascade.^22^ The third, cg03636183, is a FEV_1_/FVC ratio-associated DMS in *F2RL3*, which has been previously reported to be associated with respiratory function.^3^ This DMS was also the most informative DMS in the prediction model for COPD in an independent GS:SFHS prediction dataset, and also in the LBC1936 replication data.

As discussed above, *SOCS3* and *F2RL3* were found to provide discriminatory power in the prediction analysis. The inclusion of three other DMSs in *RALGDS*, *JADE1* and *SIAH2* improved the prediction of incident COPD risk in the GS:SFHS. *JADE1* is a negative regulator of Wnt signalling, which has been linked to the pathogenesis and progression of COPD.^23^ *RALGDS* is a Ras effector and regulates cellular processes such as vesicular trafficking, endocytosis, and migration.^24^ *SIAH2* upregulation mediates the ubiquitination of *NRF2* which has been previously associated with respiratory function^25^ and COPD.^26^

Functional annotation of differently methylated genes identified enrichment of the molecular function alternative splicing (appendix p43). This finding is consistent with earlier reports that genes associated with COPD (unlike those associated with other complex traits examined) have greater transcriptional complexity due to a disproportionately high level of alternative splicing.^27^ In addition, many such genes are spliced differently in COPD patients and controls.^27^

Functional analysis also identified two cellular pathways (appendix p43). JAK-STAT signalling (appendix p44) was highlighted in the EWAS data at both the DMS level and in the gene ontology analysis. Inclusion of a *SOCS3* DMS in the model improved the prediction of COPD in independent datasets. Finally, one of the modules comprising genes that showed both differential methylation with FEV_1_ in this study, and COPD-based gene expression in lung tissue^16^ was enriched for JAK-STAT signalling genes. Thus our data add further support to previous evidence^28^ for the importance of this pathway in respiratory function and COPD. Finally, axon guidance signalling (appendix p45) was highlighted by three of our analyses, adding support to the neuropathology hypothesis of COPD.^29^

To investigate the potential clinical implications of our findings we assessed the predictive properties of DMSs in the prognosis of COPD. The inclusion of DMSs provided added clinical value to established clinical variables in both the discovery and replication datasets. Clinical studies are needed to provide formal proof that changes in DNA methylation at these sites contribute causally to the pathogenesis, and can impact prognosis of COPD.

There are two main limitations to this study. Firstly, DNA methylation was quantified in peripheral blood. There is ongoing debate about whether DNA methylation from peripheral blood can serve as a surrogate marker for DNA methylation in lung tissue.^30^ The overlap between our findings and previous studies performed in lung tissue (appendix p8-9) suggest that, for at least some loci, the study of DNA methylation in blood may yield mechanistic insights. Moreover, our data demonstrate that DMSs from peripheral blood have both predictive and clinical value. As such, blood may be an appropriate tissue for the development of biomarkers, as it is easily and repeatedly accessible.

The second limitation is that DNA methylation was measured in blood samples collected at the same time that spirometry tests were performed. As such, our reported associations are subject to reverse causality. However, the integrated alterations in DNA methylation in this study and gene expression profiles in COPD^16^ and prospective predictive value of the selected DMSs provide indications that the DNA methylation alterations observed in blood may play a causal role in respiratory function. Nevertheless, longitudinal studies, with serial measurements of DNA methylation will be required to address causality formally.

In conclusion, using a large dataset and a robust methodological approach, we have identified DMSs associated with respiratory function and COPD, provided new mechanistic insights and supported previous hypotheses into impaired respiratory function and the pathogenesis of COPD. We also demonstrated that DMSs can be incorporated into existing models for predicting COPD risk, yielding better prediction than established clinical variables alone.

## Funding & Acknowledgements

This study was supported by a Wellcome Trust Strategic Award “Stratifying Resilience and Depression Longitudinally” (STRADL) (Reference 104036/Z/14/Z) and by the Sackler Foundation. Generation Scotland received core support from the Chief Scientist Office of the Scottish Government Health Directorates [CZD/16/6] and the Scottish Funding Council [HR03006]. Genotyping of the GS:SFHS samples was carried out by the Genetics Core Laboratory at the Wellcome Trust Clinical Research Facility, Edinburgh, Scotland and was funded by the Medical Research Council UK and the Wellcome Trust (Wellcome Trust Strategic Award (STRADL; Reference as above). The Lothian Birth Cohort 1936 is supported by Age UK (Disconnected Mind programme) and the Medical Research Council (MR/M01311/1). Methylation typing was supported by Centre for Cognitive Ageing and Cognitive Epidemiology (Pilot Fund award), Age UK, The Wellcome Trust Institutional Strategic Support Fund, The University of Edinburgh, and The University of Queensland. The LBC1936 work was conducted in the Centre for Cognitive Ageing and Cognitive Epidemiology, which is supported by the Medical Research Council and Biotechnology and Biological Sciences Research Council (MR/K026992/1). HCW is supported by a JMAS SIM fellowship from the Royal College of Physicians of Edinburgh. None of the study sponsors had a role in collection, analysis, interpretation of the data, in the writing of the manuscript or in the decision to submit the manuscript for publication.

